# Control effectiveness of APL formulation against dengue- and Zika-transmitting *Aedes* mosquitoes in Gia Lai province, Vietnam

**DOI:** 10.1101/2021.12.22.473821

**Authors:** Phung Thi Kim Hue, Le Tri Vien, Dau Minh Nga, Le Van Truong, Hoang Ha, Pham Thi Khoa, Le Thi Nhung, Ho Viet Hieu, Le Dung Sy, Trieu Nguyen Trung, Than Trong Quang, Tran Van Loc

## Abstract

**Background:** Dengue fever and Zika are two of the *Aedes*-borne diseases. Despite being widely used, synthetic mosquitocides become abortive for the mosquito control due to growing resistance and environmental pollution. In Gia Lai province (dengue-endemic area), a huge amount of cashew nut shell waste with roughly 100,000 tons/year has been disposed of into the environment, potentiating a high risk of pollution.

**Methodology/Principal findings:** To utilize it, anacardic acid was extracted and combined it with ethanol extract of the local lime peel, which contains limonene, to generate APL formulation. APL robustly exhibited inhibition of egg hatching, larvicidal effect, and repellent effect against female mosquitoes from oviposition sites in the laboratory and field. The results showed that, at a dose of 12.5 ppm, the APL formulation after 24 hours of treatment demonstrated oviposition deterrence against *Ae. aegypti* (43.6%) and *Ae. albopictus* (59.6%); inhibited egg hatching of *Ae. aegypti* (49.6%) and *Ae. albopictus* (59.6%); caused larval lethality in *Ae. aegypti* (LC_50_ = 9.5 ppm, LC_90_ = 21 ppm) and *Ae. albopictus* (LC_50_ = 7.6 ppm, LC_90_ = 18 ppm). Under natural field conditions, it showed a 100% reduction in larval density after 48 and 72 hours of the APL treatment at a tested concentration of 120 mg a.i./m^2^ and maintained a mortality rate of 100% in the next 14 days.

**Conclusions/Significance:** The APL formulation is promisingly to become an environmentally friendly and highly effective biological product for future management programs of dengue and Zika-transmitting vectors. Here offer prospects in controlling critical illnesses transmitted by several mosquito species in dengue-endemic areas.

**Author summary:** The use of synthetic insecticide to control the dengue and Zika vector population has contributed to drug resistance and caused negative impacts on the environment. The plant-based insecticide should be beneficial for mosquito management in the current situation. Gia Lai province in Vietnam is a dengue-endemic area. A large amount of cashew nut shell waste gets discarded in the area every year, which imposes an increased risk of pollution. The authors took advantage of this by extracting anacardic acid to combine with ethanol extract of local lime peel (containing limonene) to produce APL formulation. This formulation demonstrated potential activities and efficiency in controlling mosquitoes transmitting disease. In the laboratory condition, at a low dose of 12.5 ppm, APL showed activities in inhibiting egg hatching, larviciding, and repelling female *Aedes aegypti* and *Aedes albopictus*. In the field condition, APL at a dose of 120 mg a.i./m^2^ thoroughly reduced the dengue larval density after two days of contact, and this effect lasted to the next 14 days. APL is a promising and environmentally friendly larvicidal product that is highly effective in controlling dengue and Zika vectors and can play as an alternative measure for vector-borne dengue in the locality.

## Introduction

Currently, approximately 2.5 billion people live in dengue-endemic areas worldwide, with 50-100 million infected people, and an average mortality rate is 2.5% annually [1]. In Vietnam, nearly 70 million people reside in areas with prevailing dengue fever prevails. With 63 cities and provinces having dengue patients, the risk of emerging a widespread outbreak is very high [2]. According to the statistical data of the Vietnam General Department of Preventive Medicine in 2019, the number of more than 200,000 dengue incidents was three times higher than in the same period in 2018, with 50 deaths. In 2020, the entire country reported 65,046 infected cases and seven deaths. Gia Lai is one of the five provinces in the Central Highlands where the incidence of dengue fever is relatively complicated. In 2017, the number of reported dengue fever cases increased in Pleiku city (782), Chu Puh district (586), and others. Although dengue fever in Gia Lai is on a downward trend, the risk of a disease outbreak remains very high since the seasonal cycle of dengue fever is associated with the growth of the disease-transmitting vector population [3]. That seasonal occurence is four years in Northern Vietnam and three years in the Central Highlands of Vietnam. Furthermore, dengue fever in the region is affected by weather factors (often occurring in the wet season), along with habits of water storage and poor environmental sanitation (e.g., neglected containers and domestic waste trapping standing water and livestock-rearing areas with inadequate water systems and sanitation), which have facilitated various favorable breeding sites for the procreation of *Aedes aegypti* and *Aedes albopictus*, and the emergence of the outbreak [4]. Meanwhile, the misuse of insecticides floating on the market in the living space of the locals is a matter of great concern [5]. Although an effective vaccine is yet to be found, insect surveillance and vector control remain the essential solutions to prevent the spread of dengue fever [6].

So far, one of the methods for vector control against the *Aedes* mosquito is the synthetic insecticide; e.g., organophosphate and pyrethroid are popular chemical means to battle the mosquito population worldwide [7]. While *Ae. aegypti* and *Ae. albopictus* have distinctive biological, population and behavioral characteristics, their distributions continuously change throughout time, region, and ecological contexts, demanding measures and prevention strategies against these two mosquito species that have unique peculiarities to be effective [8]. Moreover, mosquito control programs have been unsuccessful due to the increasing number of mosquitoes developing drug resistance [9–11]. In Vietnam, *Ae. aegypti* became resistant to DDT, Permethrin, and Deltamethrin in the Southern area and Central Highlands [12]; it developed chemical resistance to pyrethroid group in the Southern Vietnam [13] and Hanoi [14]. Many organochlorides and organophosphates harmfully impact on biological systems and the environment [15].

The unfavorable effects of synthetic chemicals have fueled the introduction of alternative environmentally-friendly solutions to manage the mosquito. Many studies indicated that plant-based extract and essential oil had the potential to control mosquitoes [16,17]; tobacco extract of *Sphaeranthus indicus* showed potency against *Ae. aegypti* by preventing ovulation and expressing the ovicidal effect [18]. Orange extract (*Citrus sinensis*) could eliminate *Aedes* larvae with LC_50_ of 52.27 ppm [19]. Liebman supposed that different populations could express distinct drug resistance requiring tailored chemicals to control [10]. For that reason, we are concerned *Anacardium occidentale* (*Anacardiaceae*), which is widely grown in Vietnam [20]. Recent studies have suggested that the active ingredients of the cashew plant are likely to be used as an active larvicide [21]. Cashew nut shell liquid that contains phenols could kill mosquito larvae [22]. As it was extracted by solvent, its main ingredients included anacardic acid (62.9%), cardol (23.98%), and cardanol (6.99%) that was possible to eliminate *Aedes* larvae [23].

In this study, we extracted anacardic acid, an active ingredient of cashew nut shell, to combine with lime peel extract containing limonene to create APL formulation, with the purpose of producing a product having high effectiveness, exerting ovicidal and larvicidal effects against *Aedes* mosquitoes, and deterring mosquitoes from laying eggs in water. In the midst of the complicated endemic of dengue fever, dengue-transmitting mosquito control using biological product is essential to prevent mosquito-borne illnesses for public health promotion in this locality.

## Materials and methods

### Preparation of the APL formulation

From cashew nut shell waste (*Anacardium occidentale L*.) collected in Mang Yang district in Gia Lai province in Vietnam, 10 kg grinding with 50 liters of *n*-hexane was placed in a 100-liter extractor with stirring and reflux condenser and heated at 50°C for 3 hours. The extraction liquid was filtered, and the residue was extracted for the second time with 50 liters of *n*-hexane under the same above condition. The residue was processed for the third extraction with 50 liters of 95% ethanol. The solvent was distilled in vacuum at 50°C to obtain 3.5 kg of extract.

3.5 kg of cashew nut shell extract was diluted in 18 liters of acetone in a 50-liter round-bottom flask and stirred with a magnetic stirrer. The sample was slowly added 1200 grams of calcium hydroxide at room temperature. After finishing, the reaction mixture was turned up to 50°C and stirred steadily for 4 hours. After the reaction cooled down to room temperature, the precipitated calcium anacardate was filtered and washed with acetone (600 mL x 2 times), then dried to obtain calcium anacardate. 1350 grams of calcium anacardate were dissolved in distilled water (1200 mL). Then 2400 mL of HCl 11N was added and stirred continuously for 30 minutes. The sample was extracted by *n*-hexane (900 mL x 2 times). The extracting liquid *n*-hexane was washed with distilled water (600 mL x 2 times), anhydrous with Na_2_SO_4_, distilled the solvent to obtain 1.5 kg of anacardic acid.

5 kg of lime peel powder was loaded into a 100-liter round-bottom flask, added with 50 liters of 95% ethanol, heated at 40°C for 3 hours, filtered and distilled the solvent to receive the first extract. The obtained residue was processed twice more extractions under the same conditions as above, combined to receive 201 grams of lime peel extract.

The APL formulation used to test controlling activities in relation to the *Aedes* mosquito stages in the form of powder tablet was composed of anacardic acid, lime peel ethanol extract, and additives in a 3:1:6 proportion. The obtained anacardic acid was dissolved in ethanol, mixed with additives, including soluble starch and soluble sugar, and added with ethanol-dissolving lime peel extract and stirred well. Later, the mixture sample was dried with cold air and compressed to form tablets of APL formulation. This formulation was evaluated for ovicidal, larvicidal, and adult mosquito repellent effects against *Ae. albopictus* and *Ae. aegypti* in the laboratory and field conditions in Gia Lai province, Vietnam.

### Life stages of *Ae. aegypti* and *Ae. albopictus* in the study

The life stages of the mosquitoes in this study were taken from the laboratory (LAB) of the Institute for Global Health Innovations at Duy Tan University; they were not exposed to pathogens, insecticides, or repellents (LAB strains). At the same time, larvae of *Ae. aegypti* and *Ae. albopictus*, which were caught in the field (field strain) in Chu Puh district and Pleiku city in Gia Lai province through surveys and monitoring, were reared in LAB under the following conditions: temperature (T) = 28 ± 1°C, relative humidity (RH) = 70-75%, and optical cycle = 11 ± 0.5 hours.

### Examination of oviposition deterrence of APL formulation

The oviposition deterrent effect was evaluated by applying the method of Rajkumar [24] with appropriate modifications. Female mosquitoes of *Ae. aegypti* and *Ae. albopictus* (10 days old and 2 days after blood feeding) were transferred to separate cages (45 cm x 45 cm x 45 cm) made of mosquito net and muslin doors in the front for easy access. In each cage, plastic cups contained 200 mL of water placed at opposite corners. One cup was processed by one concentration of APL formulation, one cup was used as a positive control (Azadirachtin, a plant-based insecticide), and the other cup was a negative control. The tested concentrations were 6.25 ppm, 12.5 ppm, 25 ppm, and 50 ppm. Each concentration was repeated in triplicate. A sucrose solution (10%) was given to feed mosquitoes in the cages during the study period. The experiments were carried out under the conditions (T: 28 ± 1°C; RH: 70–75%) for 72 hours. After 72 hours, the eggs were counted under the microscope (Su *et al*., 1998), and the egg number laid in each cup was recorded.

The percentage of oviposition repellence for each concentration was calculated using the following formula:

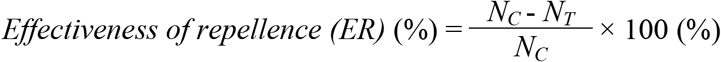

where, *N_C_* is the number of eggs in the control group and *N_T_* is the number of eggs in the experimental group.

### Examination of egg hatch inhibition of APL product

The ovicidal activity of the *Aedes* mosquito was studied according to the Elango method with appropriate adjustments [25]. New eggs laid by female *Ae. aegypti* and *Ae. albopictus* were collected and placed on filter paper liner, counted, and put into egg cups corresponding to each species. Each cup contained 25 eggs individually exposed to four different concentrations of APL formulation (6.25 ppm, 12.5 ppm, 25 ppm, and 50 ppm). Each cup was maintained separately and each concentration was repeated five times. A cup of dechlorinated tap water was served as a negative control. Azadirachtin was used as a positive control for comparison with five replicates. After 120 hours of treatment, the eggs were sieved through a muslin cloth, washed thoroughly with water, and left in plastic cups, and the eggs were counted under the microscope to assess lethality. The inhibitory activity of egg hatching was evaluated with the following formula:

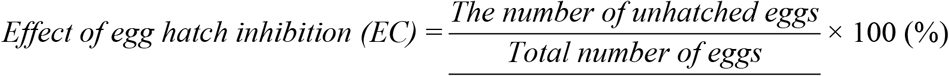

### Laboratory testing of the larvicidal effect of APL formulation against *Aedes* larvae

Biological tests for the larvicidal effect of the APL formulation against *Aedes* larvae were performed according to World Health Organization [26] bioassay protocol, with modifications, to determine the lethality concentration for mortality of 50% (LC_50_) and 90% (LC_90_). Four concentrations of the APL formulation were applied, including 6.25 ppm, 12.5 ppm, 25 ppm, and 50 ppm, each of which was repeated four times. Six plastic cups containing 249 mL of sterilized water were used, in which each cup contains 25 third and fourth instar larvae for each species (*Ae. albopictus* and *Ae. aegypti*). A beaker of (dechlorinated) tap water served as a negative control. Temephos served as a positive control. The lethality rate was observed every 24 hours until 72 hours after the experiment onset. LC_50_ and LC_90_ values were calculated by statistical analysis using a computer program.

Dead larvae are immovable larvae, and those that have gone through the pupation process will be excluded. If the proportion of pupated larvae is more than 10% in the course of the experiment, the examination will be aborted and redone. If the mortality rate in the control sample ranges from 5% to 20%, the mortality of the APL-treated groups needs to be adjusted according to Abbott’s formula (3):

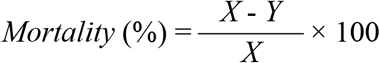

Where, *X*: percentage survival in the untreated control; *Y*: percentage survival in the treated group.

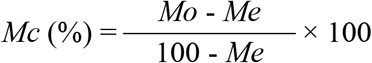

Where, *Mo*: mortality rate observed in the treated sample (%), *Me*: mortality rate observed in the control sample (%), and *Mc*: adjusted mortality rate (%). All experiments were carried out in LAB at a temperature of 28°C ± 0.5°C.

### Field testing of larvicidal activities of APL formulation

Field evaluation of the larvicidal activity of the APL formulation was conducted in Chu Puh district (Nhon Hoa town, Ia Hru commune) and Pleiku city (Yen Do ward and Yen The ward) of Gia Lai province, Vietnam. Initially, these locations were inspected to determine the suitability of different breeding grounds (water containers) of the target mosquito species for the examination. The larvicidal effect was performed on the late third instar and early fourth instar larvae of *Ae. albopictus* and *Ae. aegypti* in the field setting.

The density of larvae was measured by using a standard larvae dipper (capacity: 300 mL and diameter: 9 cm). A tested dose of 120 mg a.i./m^2^ was identified based on evaluation results in LAB conducted on *Ae. albopictus* and *Ae. aegypti*. Five kg of APL was applied to a surface area of 30 m^2^ at different breeding sites of *Ae. albopictus* and *Ae. aegypti*.

Temephos at a dose of 120 mg a.i./m^2^ was used as a positive control, and the control group was the untreated batch. Larval densities were recorded one day before application to the experimental and control water containers. The observations were then recorded at 24, 48 and 72 hours after treatment. Additional observations were made up to the end of 2 weeks. The percentage of reduction was calculated for the third and fourth instar larvae using the formula:

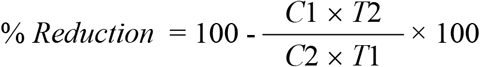

Where, *C*1, *C*2 are the larval density before and after treatment in the control group; *T*1, *T*2 are the larval density before and after treatment in the experimental groups, respectively.

### Statistical analysis

The Probit’s analytical method [27] was used to determine LC_50_ and LC_90_; corresponding confidence intervals were calculated using Statplus software. Data were analyzed for variance and compared by Tukey’s test (P < 0.05).

## Results

### Preparation of APL formulation

With the methods built the above, from the cashew nut shell source collected in Mang Yang district, Gia Lai province, Vietnam, the obtained concentrated extract achieved an efficiency of 35%. GC-MS analysis showed that it contained 41.59 % anacardic acid compared to the total obtained extract (Fig 1).

**Fig 1.**
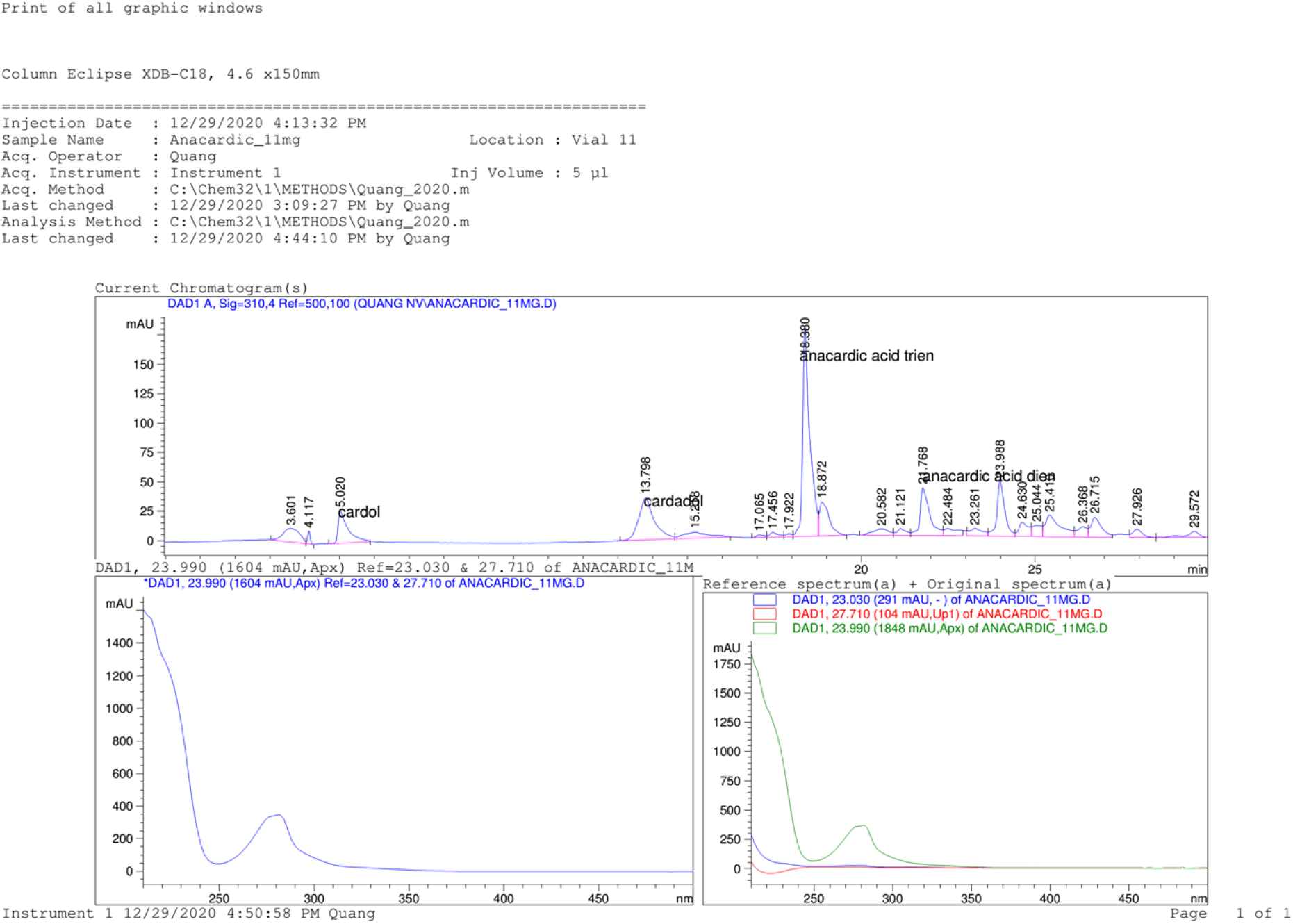
Chromatography analysis (GC-MS) of hexane+ethanol extract composition from cashew nut shell.

From the extract obtained, anacardic acid was separated with an efficiency of 42.8% compared to the crude extract and an efficiency of 15% compared to the original nut shell (Figs 2 and 3).

**Fig 2.**
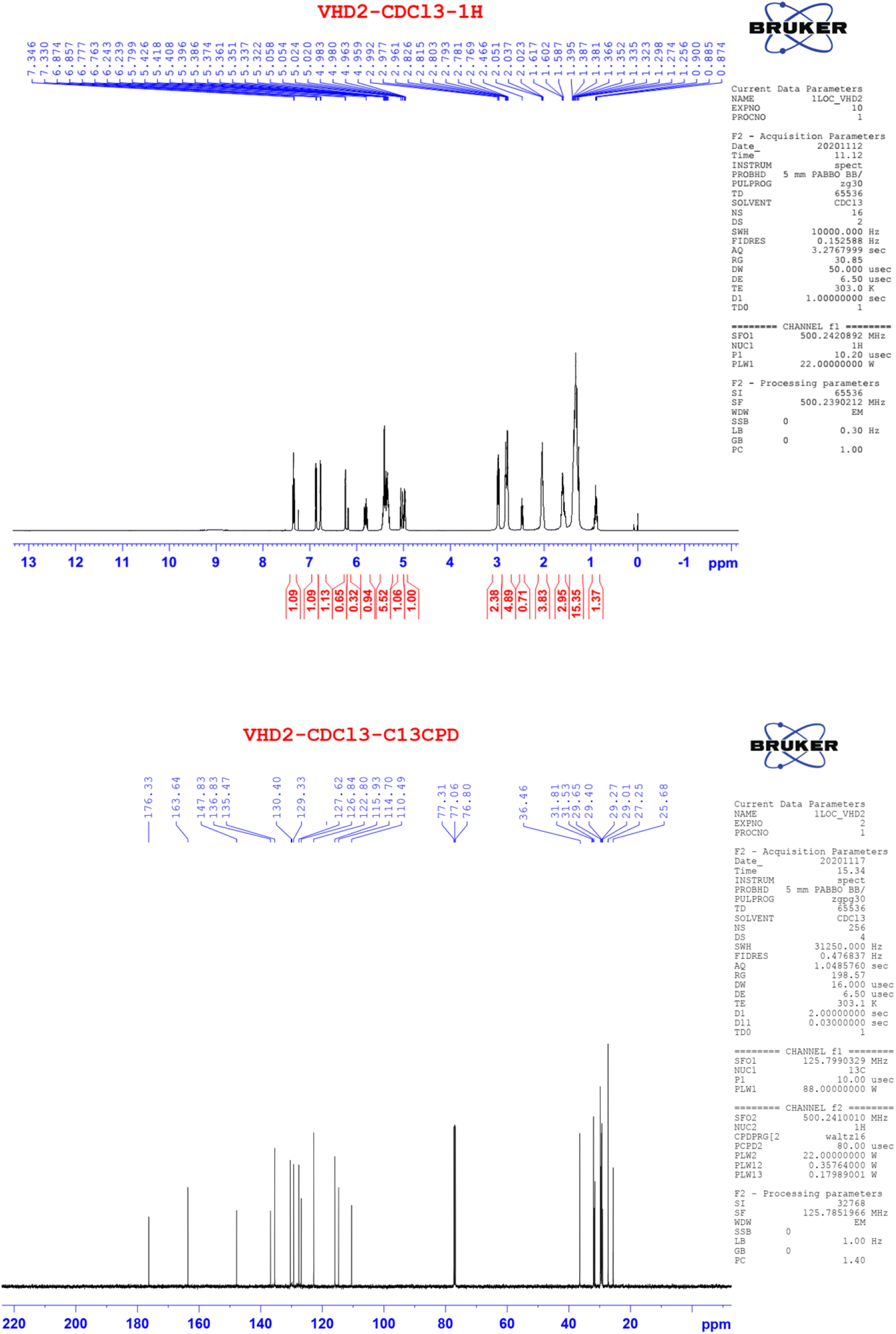
Representative total proton (^1^H) (upper) and carbon (^13^C) spectrum (lower) of anacardic acid isolated from cashew nut shell extract.

**Fig 3.**
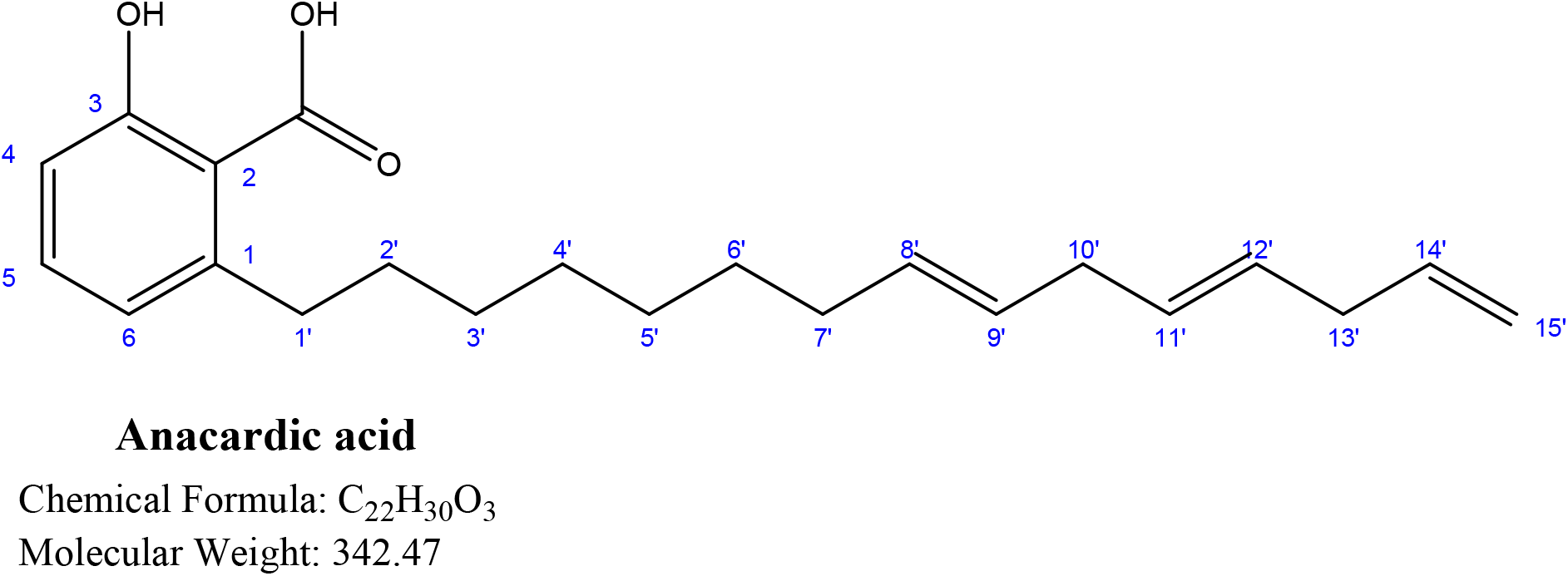
Structure of anacardic acid isolated from cashew nut shell extract.

APL formulation contained 30% anacardic acid, 10% lime peel extract, 50% soluble starch and 10% soluble sugar. Each tablet weighed 50 mg.

### Mosquito control effectiveness of APL formulation against *Aedes* mosquitoes

#### Oviposition deterrent effects of APL formulation at water breeding sites

APL showed the inhibitory activity of reproduction at all concentrations tested against adult females of *Ae. albopictus* and *Ae. aegypti* (Tables 1 and 2).

**Table 1.** Percentage of reproductive suppression in treated samples against adult female *Aedes aegypti*.

| Sample | 6.25 ppm | 12.5 ppm | 25 ppm | 50 ppm |
| --- | --- | --- | --- | --- |
| APL | 39.1 ± 1.79 | 43.6* ± 2.09 | 56.4* ± 1.15 | 68.5* ± 1.37 |
| Negative control | 1.66 ± 2.19 | 0.81 ± 1.78 | 0.80 ± 1.77 | 0.65 ± 1.75 |
| Lime peel extract | 6.50 ± 2.45 | 25.7 ± 2.19 | 40.5* ± 1.09 | 58.9* ± 1.90 |
| Anacardic acid | 3.69 ± 2.11 | 4.11 ± 1.08 | 8.80 ± 1.75 | 9.50 ± 1.15 |
| Azadirachtin | 52.1* ± 1.14 | 59.4* ± 1.09 | 63.6* ± 4.35 | 70.5* ± 1.39 |

Mean ± Standard deviation (SD) (n=5); ppm = parts per million; Statistical significance: \*, $p \leq 0.05$ and \*\*, $p \leq 0.01$

**Table 2.** Percentage of reproductive suppression in treated samples against adult female *Aedes albopictus*.

| Sample | 6.25 ppm | 12.5 ppm | 25 ppm | 50 ppm |
| --- | --- | --- | --- | --- |
| APL | 48.8 ± 5.17 | 59.6* ± 2.97 | 69.6* ± 1.48 | 78.1** ± 1.12 |
| **Negative control** | 1.06 ± 2.19 | 0.87 ± 1.58 | 0.86 ± 1.78 | 0.75 ± 1.79 |
| **Lime peel extract** | 8.90 ± 2.70 | 39.2* ± 2.04 | 52.9* ± 3.45 | 69.6* ± 1.35 |
| **Anacardic acid** | 3.98 ± 2.19 | 4.90 ± 2.06 | 9.80 ± 3.74 | 9.90 ± 1.28 |
| **Azadirachtin** | 50.3* ± 2.14 | 61.6* ± 1.18 | 69.6* ± 4.30 | 75.6* ± 1.92 |

Mean ± Standard deviation (SD) (n=5); ppm = parts per million; Statistical significance: \*, $p \leq 0.05$ and \*\*, $p \leq 0.01$

Compared to the negative control, at a concentration of 50 ppm, the APL formulation showed the highest impact on two *Aedes* mosquito species in the laboratory, in particular, for *Ae. aegypti*, was recorded to be 68.5%, for *Ae. albopictus*, was 78.1% in oviposition deterrent activities with statistical significance.

At 12.5 ppm, Azadirachtin demonstrated oviposition deterrent activity at water breeding sites of 59.4% (*Ae. aegypti*) and 61.6% (*Ae. albopictus*). This activity of Azadirachtin against *Ae. aegypti* and *Ae. albopictus* was higher than that of the APL formulation at every concentration, but the difference was not considerable between each concentration (P > 0.05). At 50 ppm, lime peel extract showed inhibition of oviposition of 69.6% against *Ae. albopictus* and 58.9% against *Ae. aegypti*, while the impact of anacardic acid was only 9.9% (*Ae. albopictus*) and 9.5% (*Ae. aegypti*). The APL containing anacardic acid and lime peel extract had activity approaching 78.1% (*Ae. albopictus*) and 68.5% (*Ae. aegypti*).

### Inhibition of egg hatching of APL formulation

Among different concentrations of APL, they showed the inhibitory effect on the egg hatching of *Ae. aegypti* and *Ae. albopictus* (Tables 3 and 4). APL exhibited this activity with the highest concentration of 88.6% (*Ae. albopictus*) and 78.5% (*Ae. aegypti*) at a concentration of 50 ppm compared to the negative control with statistically significance.

**Table 3.**
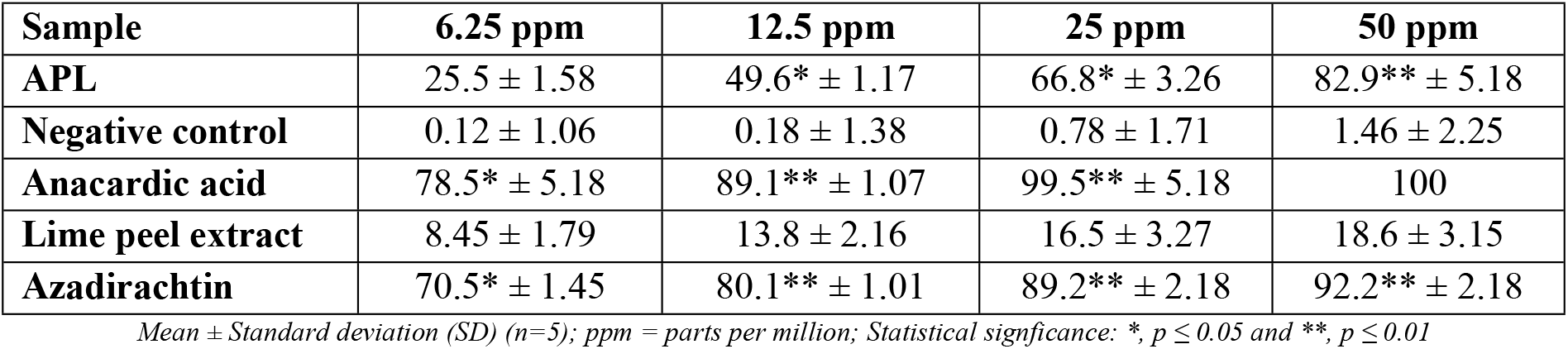
Percentage of ovicidal activity against egg hatching of *Aedes aegypti* in the samples.

**Table 4.** Percentage of ovicidal activity against the egg hatching of *Aedes albopictus* in the samples.

| Sample | 6.25 ppm | 12.5 ppm | 25 ppm | 50 ppm |
| --- | --- | --- | --- | --- |
| APL | 29.8 ± 1.27 | 59.6* ± 1.77 | 76.3* ± 1.25 | 90.6** ± 3.17 |
| Negative control | 0.15 ± 1.09 | 0.19 ± 1.35 | 0.38 ± 1.21 | 1.45 ± 2.05 |
| Anacardic acid | 80.5** ± 3.15 | 95.6** ± 1.79 | 100 | 100 |
| Lime peel extract | 10.5 ± 2.71 | 14.9 ± 3.15 | 17.8 ± 2.28 | 19.9 ± 1.18 |
| Azadirachtin | 72.5** ± 3.18 | 81.8 **± 3.18 | 90.1** ± 1.19 | 100 |

Mean ± Standard deviation (SD) (n=5); ppm = parts per million; Statistical significance: \*, $p \leq 0.05$ and \*\*, $p \leq 0.01$

At 50 ppm, anacardic acid suppressed egg hatching by 100% in both *Ae. albopictus* and *Ae. aegypti*, while the effect of lime peel extract was 19.9% (*Ae. albopictus*) and 9.5% (*Ae. aegypti*). The APL containing both ingredients presented activities of 88.6% (*Ae. albopictus*) and 82.9% (*Ae. aegypti*).

The Azadirachtin control (12.5 ppm) reported egg hatch inhibitory activity of 80.1% and 81.8% against *Ae. aegypti* and *Ae. albopictus*, respectively, the results did not have substantial differences in the potency of Anacardic acid at the same concentration. APL values at concentrations of 12.5 ppm to 50 ppm were lower than those of azadirachtin and anacardic acid, although the inhibitory effect of APL on egg hatch was statistically significant (P < 0.05) compared with negative control. The lime peel extract did not show the inhibitory effect of egg hatching against *Ae. aegypti* and *Ae. albopictus* with statistical significance.

### Laboratory evaluation results of larvicidal effect of APL formulation

The mean LC_50_ and LC_90_ values of the APL formulation against *Ae. albopictus* and *Ae. aegypti* are shown in Table 5. LC_50_ values of APL were 7.6 ppm and 9.5 ppm, while the LC_90_ values were 18 ppm and 21 ppm with respect to *Ae. albopictus* and *Ae. aegypti*, respectively. No substantial differences were observed between field strain and LAB strain (P > 0.05).

**Table 5.** Larvicidal activity of APL formulation against *Aedes* mosquitoes in the laboratory.

| Vector species | Larvicidal effect (ppm) |  |
| --- | --- | --- |
| | $LC_{50}$ | $LC_{90}$ |
| *Aedes aegypti* | 9.5 ± 0.4 | 21 ± 0.2 |
| *Aedes albopictus* | 7.6 ± 0.1 | 18 ± 0.3 |

Water depth: 2.5 cm; Mean ± SD; Confidence interval 95%; Number of repeats n = 5

The reference control (temephos) showed that the LC_50_ and LC_90_ values against the LAB strain of *Ae. albopictus* and *Ae. aegypti* were 6.6 ppm & 12 ppm and 7.9 ppm & 19 ppm, respectively. For field strains, the values of LC_50_ and LC_90_ at 24 hours of exposure were 8.5 ppm & 20 ppm and 9.5 ppm & 27 ppm, respectively, with a significant difference between field strain and LAB strain (P < 0.05).

### Field evaluation results of larvicidal effect of APL formulation

Before the onset of treatment, our preliminary survey conducted at the breeding sites of female *Aedes* mosquitoes showed that, among the 2816 water containers (tyre, barrel, bucket and others) found inside and outside of the households in the investigation locations (Ia Hru commune, Nhon Hoa township, Chu Puh district, and Yen The ward, Pleiku city, Gia Lai province, Vietnam), 1830 water containers (65%) were positive with the presence of mosquito larvae, *Ae. aegypti* and *Ae. albopictus*, and the larvae were primarily settled in old tyres.

The larvicidal activities of the APL formulation against *Ae. aegypti* in different water containers are shown in Table 6. Larval density decreased from 90.9% to 95.6% on day 1, then continued from 92.5% to 97.9% on day 2, dropped to 99.2% to 100% on day 3, and reached 100% on day 7. Among those, tyres, buckets, and barrels had a reduction rate of *Ae. aegypti* larvae >90% on the first day and continued to decrease until the seventh day by 100%. Meanwhile, the positive control, temephos at the same concentrations, has demonstrated a lower reduction rate in larval density (86.8%) in relation to the APL formulation.

**Table 6.** Larvicidal activity of APL formulation against *Aedes aegypti* in the field.

| Containers | Larval density before treatment | pH | Percentage of larval density decrease after treatment |  |  |  |  |
| --- | --- | --- | --- | --- | --- | --- | --- |
|  |  |  | Day 1 | Day 2 | Day 3 | Day 7 | Day 14 |
| Tyre | 30.5 ± 3.6 | 8.0–9.0 | 92.5 ± 2.2 | 97.6 ± 2.8 | 99.2 ± 1.4 | 100 | 100 |
| Bucket | 24.4 ± 5.5 | 8.0–8.5 | 95.6 ± 2.5 | 97.9 ± 2.4 | 100 | 100 | 100 |
| Barrel | 41.4 ± 5.0 | 8.0–8.5 | 90.9 ± 1.6 | 92.5 ± 3.5 | 99.6 ± 4.8 | 100 | 100 |
|  |  |  | **93.0 ± 2.1** | **96.0 ± 2.9** | **99.6 ± 2.1** | **100** | **100** |

Number of repeats $n = 21$ ; Mean ± SD

The reduction in larval density of *Ae. albopictus* in different breeding grounds is displayed in Table 7. For the tyre, the percentage of larval density reduction against *Ae. albopictus* was 93.8%, 98.6%, 100% on day 1, day 2, and day 3, respectively, after treatment, and continued to maintain 100% until day 7. For water-containing buckets, the reduced larval density measured on day 1, day 2, day 3, and day 7 was 92.5%, 95.9%, 100% and 100%, respectively. In water barrels, the larval density continued to drop on day 1 (98.1%) until day 7 (100%).

**Table 7.**
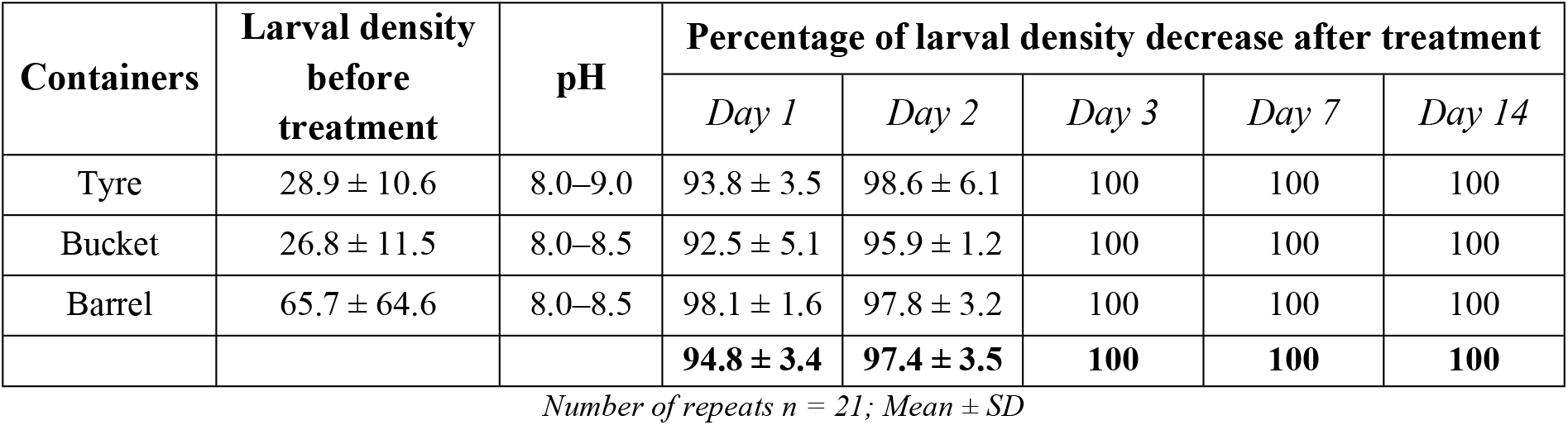
Larvicidal activity of APL formulation against *Aedes albopictus* in the field.

## Discussion

Cashews are found in many regions of Vietnam, but also here, most cashew nut shells have been discarded with unexploited potentials. While many studies reported different effects on this type of plant residue, for example, ethanol and hexane extracts of cashew nut shell showed that LC_50_ and LC_90_ of larvicidal activities against *Aedes* larvae were 3.29 mg/L & 8.13 mg/L, and 7.31 mg/L & 13.55 mg/L, respectively [28]. The cashew nut shell contains 90% anacardic acid [29], which had insecticidal effect [30], inflicted irreversible damage to the middle stage of the third instar larvae of *Ae. aegypti* [31], caused injured cytoplasm, hypertrophy of the nucleus and epithelial cell in *Aedes* larvae [31), and revealed larvicidal effect as an acetylcholinesterase inhibitor [30]. Hexane fraction of cashew nut shell (containing primarily anacardic acid) showed the highest larvicidal activities with LC_50_ of 4.01 mg/L and LC_90_ of 11.29 mg/L compared to the commercial larvicide [28]. Anacardic acid in cashew methanol extract was larvicidal in both *Ae. albopictus* and *Ae. aegypti* species [32]. Notwithstanding the potential values, about 100,000 tons of cashew nut shells have been disposed of wastefully each year in Gia Lai, Vietnam [33]. The problems of obtaining cashew nut shell extract to produce biological preparations for mosquito control should be of greater interest in this locality. In addition, another common regional plant is the domestic lime species (*Citrus aurantifolia*), a small herb widely grown in Vietnam. The lime peel’s essential oil contains D-limonene (70.37%) [34] with the limonene content found abundantly in the unripe peel [35]. Locals usually take the juice and discard the remaining. Limonene is a highly oxidizing secondary metabolite [36] and exhibited mosquito-killing characteristics [37]. It is an odorant and has various effects, especially mosquito repellence [38]. Our previous study has indicated that ethanol extract of lime peel exhibited a mosquito repellent effect against *Ae. aegypti* [39]. In the fumigation test, the LC_50_ of pure limonene against the insect was 10.65. For the contact toxicity test, the obtained LD_50_ value was 0.21 μL after 48 hours [40]. As such, the APL formulation containing anacardic acid (Fig 2 and S1 Appendix) may have the main effect on control of *Aedes* mosquitoes through egg hatch inhibition and larvicidal effect. Similarly, limonene in the formulation is known to be an odorant and may ward off the mosquitoes from their breeding sites.

To prove the presumption, the results showed that APL suppressed reproduction at all concentrations tested against adult female *Ae aegypti* and *Ae. albopictus* (Tables 1 and 2). At 12.5 ppm (Figs 4A and 4B), the influences of APL formulation were 43.6% on *Ae. aegypti* and 59.6% on *Ae. albopictus* (P < 0.05). At 50 ppm, the oviposition deterrence reached the highest level, i.e. 68.5% for *Ae. aegypti* and 78.1% for *Ae. albopictus* relative to the negative control with statistical significance (Figs 4A and 4B). The aforementioned APL ingredients include anacardic acid and limonene. For a better understanding of their reproductive suppression activities, we employed the same concentrations as APL. As a result, lime peel extract exhibited activities against female *Ae. aegypti* and *Ae. albopictus* of 58.9% and 69.6%, respectively, while anacardic acid-treated samples showed the lowest effect in all concentrations examined (Tables 1 and 2). It denoted that the reproductive suppression activity of APL could derive from limonene properties in lime peel extract. Our previous study also proved that lime peel extract (*Citrus aurantifolia*) collected in Gia Lai in Vietnam, which contains limonene, might repel *Ae. aegypti* with protection of 70 minutes and mosquito bite rate of 0.9% [39]. In many studies, limonenes extracted from lime and citrus peel were odorous [41], had fragrant and deterred mosquitoes [37], exerted mosquito repellent and insecticidal effects [42], and enhanced repellence properties as concentration and time exposed to limonene increased [43]. Since they were volatile, odor indicators binding to the protein receptor located on the hairs of mosquito specialized olfactory sensory neurons, when exposed to the external environment, could help them escape the scented places [44], especially in stagnant sources treated with APL.

**Fig 4.**
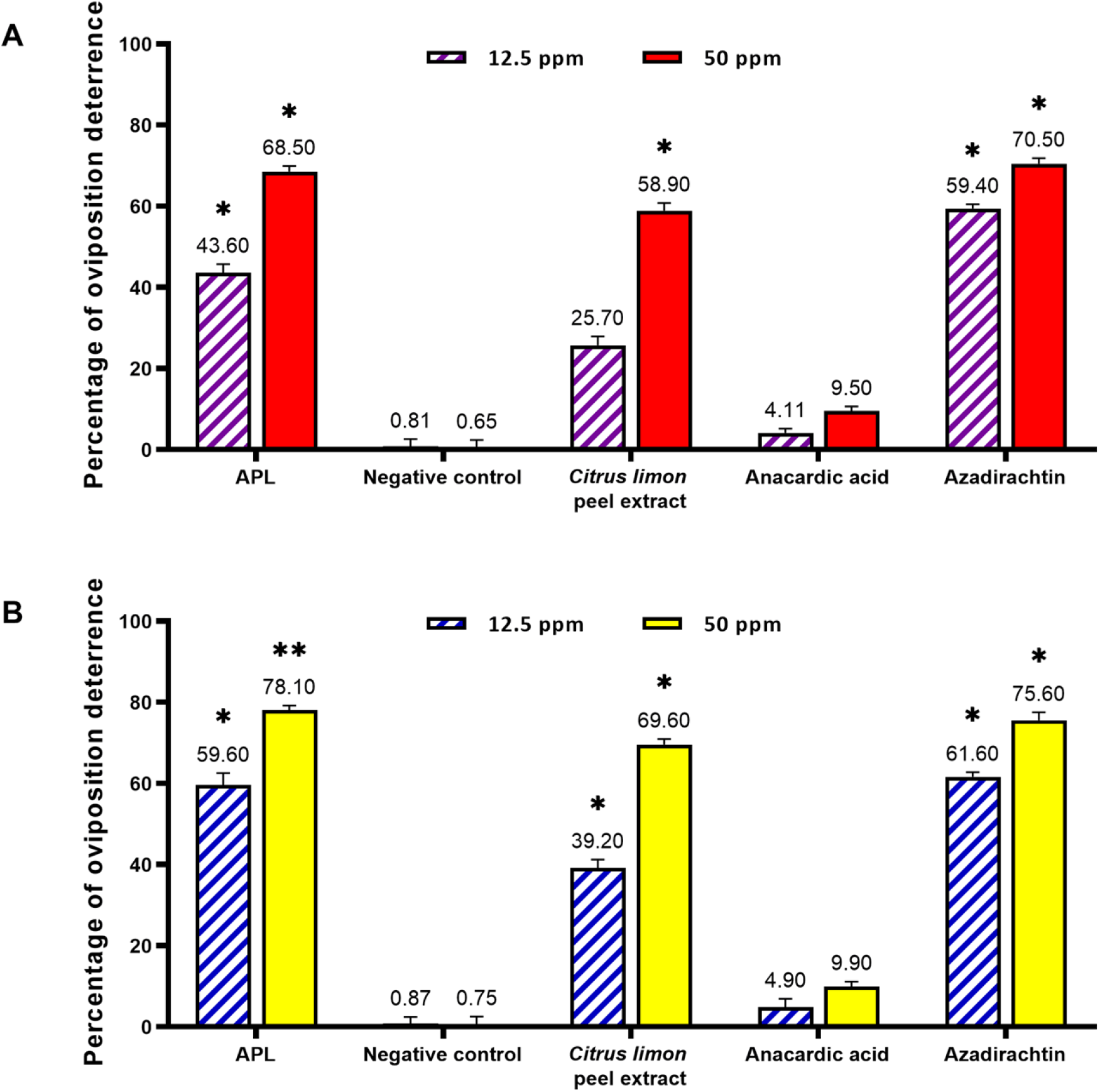
Oviposition deterrence activity of the samples against *Ae. albopictus* and *Ae. aegypti*. Oviposition deterrent effects were tested on female mosquitoes of *Ae. aegypti* (**A**) and *Ae. albopictus* (**B**) at 12.5 and 50 ppm. Results are displayed in Mean ± Standard deviation of five repeats. Abbreviation: ppm = parts per million; *, p ≤ 0.05 and **, p ≤ 0.01 denote statistical signficance.

However, the study by Alves indicated that a high proportion of anacardic acid in cashew extract, giving an oviposition activity index (OAI) of −0.1, could inhibit egg laying at the oviposition site by *Ae. aegypti* [45]. As such, the combination of anacardic acid and limonene in the APL formulation is most likely to have created a synergistic effect regarding the reproductive suppression ability in *Aedes* mosquitoes. Still, this point needs further evidence in future studies.

When treated at 12.5 ppm, Azadirachtin exhibited repellent activities that were 59.4% (*Ae. aegypti*) and 61.6% (*Ae. albopictus*) without statistically significant differences compared to APL. Azadirachtin is a plant-based insecticide accredited as the most effective product at present [46]. APL showed the oviposition inhibitory activity in two vector species with a more powerful effect than Azadirachtin (Fig 4).

Different concentrations of APL demonstrated inhibition of egg hatching upon *Ae. aegypti* and *Ae. albopictus* (Tables 3 and 4). At 12.5 ppm (Figs 5A and 5B), the positive control azadirachtin recorded the egg hatch inhibition against *Ae. aegypti* and *Ae. albopictus* were 80.1% and 81.8%, respectively. The result did not have a substantial difference (P > 0.05) compared to the effect of anacardic acid at the same concentration. The egg hatching inhibitory effect of APL was statistically significant (P < 0.05) in comparison to the negative control, but was lower than that of azadirachtin and anacardic acid. Meanwhile, ethanol extract of lime peel recorded the weakest effect against *Ae. aegypti* and *Ae. albopictus* (13.8% and 14.9%). To clarify this, we increased the concentrations of the treatment. At 50 ppm (Figs 5A and 5B), both anacardic acid and azadirachtin achieved a 100% efficiency on both species. APL also responded with an increased egg hatch inhibition of 82.9% (*Ae. aegypti*) and 90.6% (*Ae. albopictus*), while lime peel extract recorded the inhibitory effect at levels of 18.6% and 19.9% against *Ae. aegypti* and *Ae. albopictus*, respectively. These results proved that the egg hatching inhibitory effect of APL could be highly dependent on the anacardic acid present in the formulation. Farias reported that sodium anacardate resisted egg hatching in *Ae. aegypti* with an EC_50_ of 162.93 ± 29.93 mg/ml [47]. Egg hatching inhibitory activities of the cashew nut shell liquid extract against *Ae. aegypti*, which contains anacardic acid, showed LC_50_ of 8.06 mg/L and LC_90_ of 15.53 mg/L [28]. Nevertheless, Hebeish suggested that the mosquito repellent effect improved as concentration and duration of exposure to limonene increased [43]. Limonene, an active component found in essential oils, disrupted the wax surface of the insect respiratory system leading to suffocation [48]. Moreover, limonene also possessed high toxicity to insects [49]. Therefore, a small proportion of limonene in APL may only ward off mosquitoes, but not confer an ovicidal effect, which met the objective of using limonene as a flavoring agent for APL formulation.

**Fig 5.**
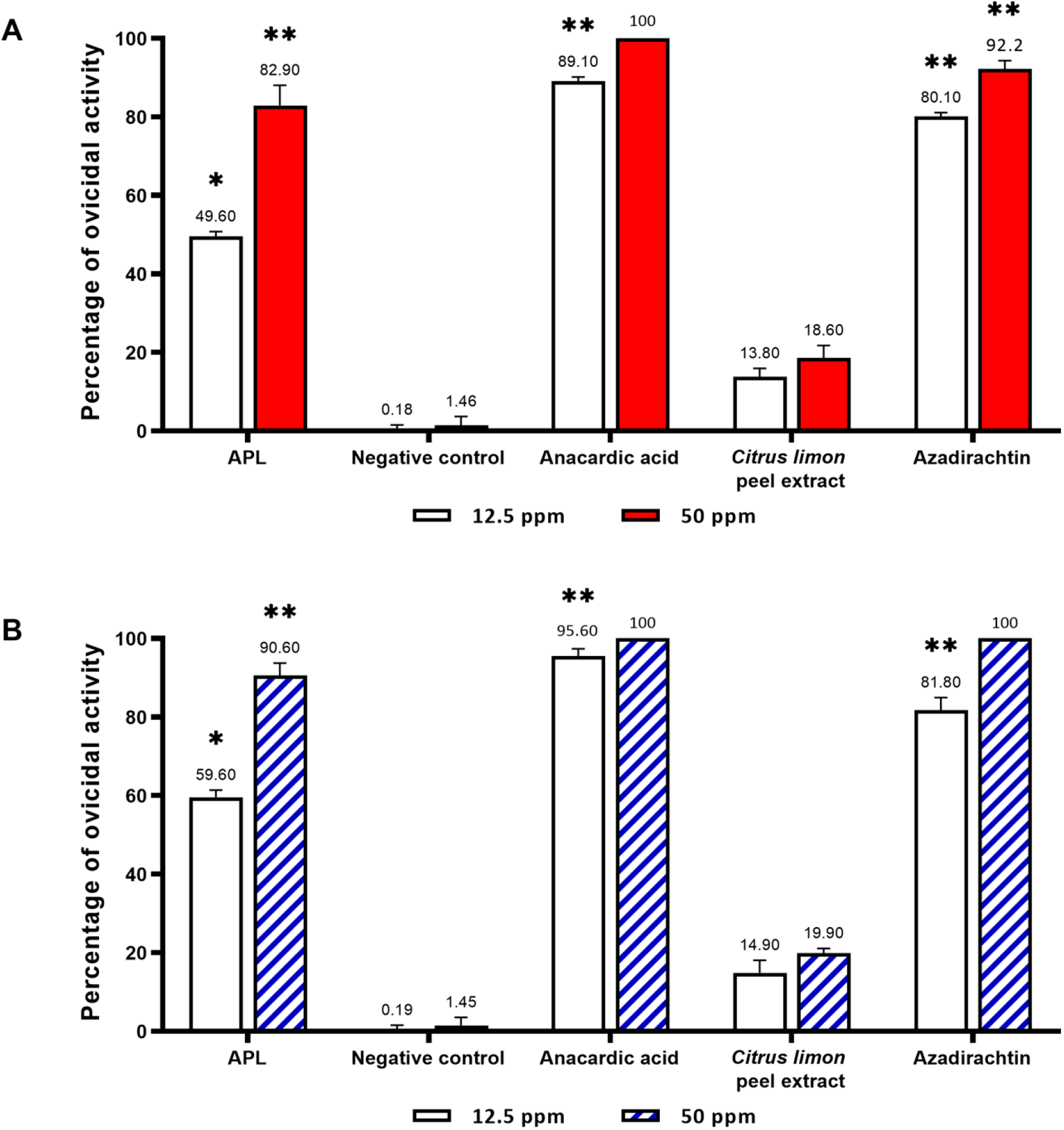
Egg-hatching inhibitory activity of the samples against *Ae. albopictus* and *Ae. aegypti*. Egg hatch inhibition was examined on female *Ae. aegypti* (**A**) and *Ae. albopictus* (**B**) at tested concentrations of 12.5 ppm and 50 ppm. Results are displayed in Mean ± Standard deviation of five repeats. Abbreviation: ppm = parts per million; *, p ≤ 0.05 and **, p ≤ 0.01 denote statistical signficance.

The evaluation of the larvicidal effect of the APL formulation revealed that the LC_50_ values were 7.6 ppm and 9.5 ppm, while the LC_90_ values were 12 ppm and 15 ppm against *Ae. albopictus* and *Ae. aegypti*, respectively (Table 5). These results showed that the APL formulation was more potent than the combination of cashew nut shell liquid and castor oil, which produced 100% lethality in larvae at a dose of 5 mg/mL [50].

The results noted no significant difference observed between the field strain and LAB strain (P > 0.05). It showed that the tested APL formulation was prospective for the larvicidal effect and demonstrated comparable results with respect to repellent activities against *Aedes* mosquitoes of both the LAB strain and the field strain. In this regard, we speculated that APL is yet to entail any resistance against the field strains, *Ae. albopictus* and *Ae. aegypti*. Meanwhile, the reference control (organophosphate larvicide temephos) exhibited values of LC_50_ and LC_90_ against two LAB strains, *Ae. albopictus* & *Ae. aegypti*, were 6.6 & 12 ppm, and 7.9 & 16 ppm, respectively, and against two field strains were 8.5 & 16 ppm, and 9.2 & 20 ppm, respectively. At the 24 hours of exposure, the LAB strains and the field strains showed significant differences (P < 0.05). These results suggested that the LAB strains were more sensitive to temephos than the field strains, which seemingly meant that *Ae. aegypti* and *Ae. albopictus* have developed resistance to synthetic larvicides. Our previous report indicated that, in the same investigation field where locals tend to abuse the commonly commercial larvicide on the market (66.11%) to remove dengue mosquitoes within their residence and the surrounding environment [5]. Therefore, it may contribute to the development of synthetic chemical resistance against *Aedes* mosquitoes in these places. Several studies have addressed similar problems in Vietnam and around the world. The laboratory strain of *Ae. aegypti* was found to be more sensitive than the field strain in response to essential oils containing pyrethroids [51]. Drug-resistant mosquitoes of *Ae. aegypti* were identified in California [10]; pyrethroids- and organophosphates-resistant *Ae. aegypti* mosquitoes were detected in the southern United States [11].

The LC_50_ and LC_90_ values of APL in this study also recorded that *Ae. albopictus* was more sensitive than *Ae. aegypti* in both laboratory strain and field strain (Fig 6). This result was aligned with previous studies, in which *Ae. albopictus* was reported to be more susceptible to plant extracts than *Ae. aegypti* [52]; *Ae. aegypti* exhibited higher resistance than *Ae. albopictus* due to its close proximity to human habitats, leading to higher exposure to insecticides [53]. In Vietnam, *Ae. aegypti* reported having developed resistance to Permethrin, Lambda-cyhalothrin, Deltamethrin, DDT in the southern region [54] and Hanoi [14]. Consequently, Liebman suggested the need for alternative measures to be implemented in regard with the emergence of specific pesticide resistance profiles [10]. Therefore, we applied the APL formulation that contains natural components, including anacardic acid, limonene, to kill *Aedes* larvae and further improve the chemical resistance situation for more effective control of dengue and Zika vectors in this locality.

**Fig 6.**
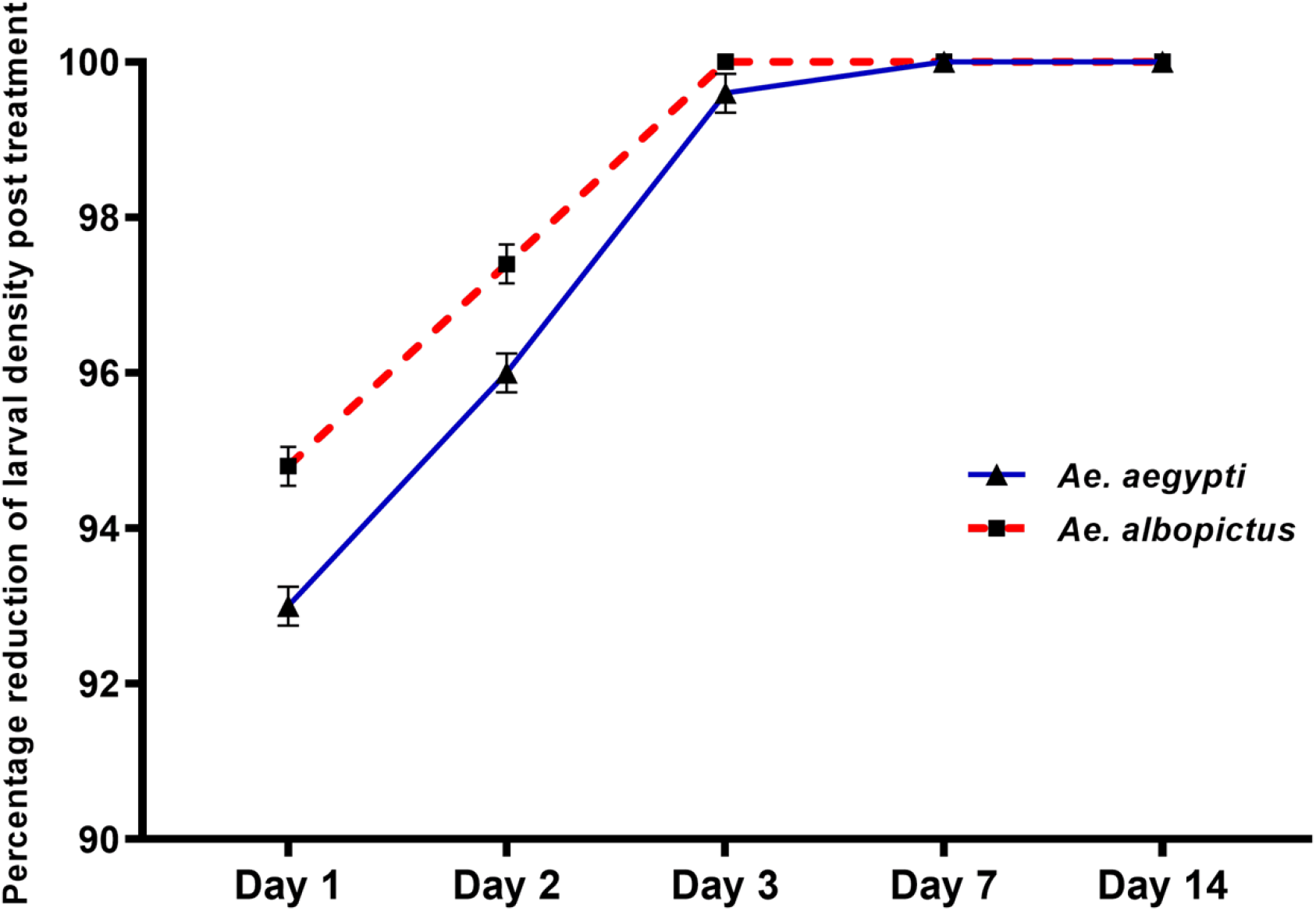
Comparison of the larvicidal efficiency of APL formulation against *Aedes albopictus* and *Aedes aegypti*. Results are displayed in Mean ± Standard deviation (SD).

Throughout the assessment, the observed larvae under the influence of APL showed signs of disintegration, unresponsiveness, and immobility, with >90% mortality rate in both *Ae. albopictus* and *Ae. aegypti* after 24-hour exposure. Histological examination revealed the disruption of intercellular junctions, possibly because the hydrophobic fatty chain of anacardic acid facilitates its penetration into the lipoprotein membrane of the larval cell and therefore affects the permeability. Scudeler *et al*. (2014) determined that the renewal of cell layers occurs as a defensive mechanism to compensate for loss from local damage induced by cellular acids. The lack of responsiveness and immobility in insects could be explained by ionic imbalance and loss of control of water intake across the plasma membrane induced under the influence of oils, toxins, and plant extracts [55]. Furthermore, the cytoplasm disruption may result from the anacardic acid effect on columnar cells [31]; and larvae immobility may be owing to inhibition of anacardic acid on acetylcholinesterase [30].

Considering the above effects, we proceeded to investigate the dosage of 120 mg a.i./m^2^ at mosquito breeding sites, including tyre, barrel, bucket, and other stagnant water, compared to control sites not treated with APL. It showed the percentage reduction of larval density by >93% in *Ae. aegypti* population on days 1, day 2, day 3; increased to 100% from day 4 to day 7; and was maintained until day 14. Meanwhile, the density of the *Ae. albopictus* population showed a >94.8% decrease from day 1 to day 2, reduced by 100% from day 3 to day 7, and lasted until day 14. The larval density reduction level exhibited a potent effect relative to the positive control, temephos (a synthetic insecticide).

In the past five decades, synthetic chemical insecticides have been employed imprudently to prevent the *Aedes* mosquito. These resulted in numerous adverse effects on human health and the environment [56], directly associated with genetic instability and increased predisposition to cancer [57,58], and caused hormonal disturbances and male infertility [59]. These downsides of synthetic chemicals have raised the necessary awareness about the demand for eco-friendly, specific-targeted, low-cost, and highly effective insecticides for vector control [60]. Furthermore, biological insecticides usually affect a small scale with rapid degradation, leading to lower exposure levels and fewer environmental contaminations [61]. However, plant-based insecticides only account for <1% in the global market [62]. For that reason, the search for novel substances, such as extracts of cashew nut shell and lime peel, has given a high likelihood of receiving special attention from the community due to their environmentally friendly attributes.

Our previous study noted that, at a concentration of 500 mg/L, cashew nut shell extract did not affect sailfish and lettuce growth [33]. The HKM formulation consisted of 1% lime peel extract and 10% cashew nutshell extracts (cardol removed) were tested to be safe in mice at a dose of 500 mg/kg body weight [63]. The available studies also revealed that cashew nut shell extract was non-toxic and could not induce genetic mutation at any concentration [64]. Pregnant female mice given a mixture of cashew nutshell extract and castor oil (TaLCC-20) at an oral dose of 5 mg/kg showed stable embryonic development, but did not show any genetic damage and side effects in mammals, including humans [50]. Sodium anacardate, isolated from the cashew nut shell, was harmless [47]. Furthermore, the cashew nutshell liquid extract containing 76.93% anacardic acid did not show any phytotoxic effect at a concentration of 3% [65]. The above studies supported the results obtained from this study. It also showed that the employment of abundant sources of plant residues, like cashew nut shell and lime peel, to create the APL formulation as a new and green vector control approach could be considered an alternative solution to tackle the spread of mosquitoes that transmit dengue and Zika virus and simultaneously resolve environmental pollution caused by plant residues and synthetic insecticides in this locality.

## Conclusion

Vector control has become a crucial issue in restraining the mosquito-borne endemic. This study has proposed APL formulation, a potential bioinsecticide alternative made from cashew nut shells and lime peel waste. This formulation is easy to produce with low technology, has the advantage of being environmentally friendly and effective, and can efficiently incorporate into existing mosquito control programs. In this way, the expense for mosquito management can reduce in place of operating conventional chemical control measures. The APL formulation showed many encouraging activities against *Ae. aegypti* and *Ae. albopictus*, including egg hatch inhibition, oviposition deterrence in the laboratory, and larvicidal effects in the laboratory and field. It is not only a green product substitute suitable for the prevention of dengue and Zika vectors but also overcomes ecological risks through biodegradable properties and demonstrates larvicidal effects to serve as a new protection measure for the stockpile of plant-based insecticides.

## Acknowledgments

We send appreciation to the contribution of the technicians who partially participated in this research from the microbiology laboratory of Institute for Global Health Innovations in Duy Tan University, technicians from the Institute of Health Research and Educational Development in Central Highlands, Science Services Of Insect Joint Stock Company, and Institute of Chemistry in Vietnam Academy of Science and Technology.

## Supporting information

**S1 Appendix. Additional information about isolation of anacardic acid from cashew nut shell**.

